# LRRK2 kinase inhibitors induce a reversible effect in the lungs of non-human primates with no measurable pulmonary deficits

**DOI:** 10.1101/390815

**Authors:** Marco A.S. Baptista, Kalpana Merchant, Ted Barrett, Diane K. Bryce, Michael Ellis, Anthony A. Estrada, Matthew J. Fell, Brian K. Fiske, Reina N. Fuji, Paul Galatsis, Anastasia G. Henry, Sue Hill, Warren Hirst, Christopher Houle, Matthew E. Kennedy, Xingrong Liu, Matthew L. Maddess, Carrie Markgraf, Hong Mei, William A. Meier, Stephen Ploch, Christopher Royer, Karin Rudolph, Alok K. Sharma, Antonia Stepan, Stefan Steyn, Craig Trost, Zhizhang Yin, Hongshi Yu, Xiang Wang, Todd B. Sherer

## Abstract

Putative gain-of-function mutations in leucine-rich repeat kinase 2 (LRRK2), resulting in increased kinase activity and cellular toxicity, are a leading genetic cause of Parkinson’s disease (PD). Hence, there is strong interest in developing LRRK2 kinase inhibitors as a disease-modifying therapy. Published reports that repeat dosing with two LRRK2 kinase inhibitors (GNE-7915 and GNE-0877) induce histopathological changes in the lung of non-human primates Fuji et al. 2015 (*1*) raised concerns about potential safety liability of LRRK2 kinase inhibitors. In the present study, we sought to determine whether previously observed effects in the lung: (a) represent on-target pharmacology, but with the potential for margin of safety, (b) are reversible upon drug withdrawal, and (c) are associated with pulmonary function deficits. To this end, we evaluated the histopathological effects, toxicokinetics and target inhibition of three structurally diverse LRRK2 kinase inhibitors, GNE-7915 (30 mg/kg, BID, as a positive control), MLi-2 (15 and 50 mg/kg, QD) and PFE-360 (3 and 6 mg/kg, QD) following 2 weeks of dosing in non-human primates. Subsets of animals dosed with GNE-7915 or MLi-2 were evaluated after 2-week dose-free periods. All three LRRK2 kinase inhibitors induced mild cytoplasmic vacuolation of type II pneumocytes, as reported previously, confirming an on-target effect of these compounds. Interestingly, despite lower doses of both PFE-360 and MLi-2 producing nearly complete inhibition of LRRK2 kinase activity in the brain as assessed by levels of pS935-LRRK2, histopathological changes in lung were absent in animals treated with low-dose PFE-360 and observed only sporadically in the low-dose MLi-2 group. The lung effect was fully reversible at 2 weeks post-dosing of GNE-7915. In a second study of identical dosing with MLi-2 and GNE-7915, no deficits were observed in a battery of translational pulmonary functional tests. In aggregate, these results do not preclude the development of LRRK2 kinase inhibitors for clinical investigation in Parkinson’s disease.

## Introduction

Parkinson’s disease (PD) is the second most common progressive and age-related neurodegenerative disorder characterized by the loss of nigrostriatal dopamine neurons and accumulation of intracellular protein aggregates termed Lewy bodies (*2*). With an aging population, the prevalence of PD worldwide and the associated health care burden is expected to increase exponentially (*3*). Currently available drugs primarily augment dopaminergic neurotransmission to treat motor symptoms of PD, but do not affect the progression of the disease. Thus, there remains an urgent need to develop disease-modifying therapies for PD.

Genetic studies have provided unprecedented insights into the pathogenesis of PD and suggested novel drug targets (*4*). One such target is leucine-rich repeat kinase 2 (LRRK2), which has attracted significant interest from the pharmaceutical industry for the following reasons: (a) point mutations in the *LRRK2* gene represent the most common causes of genetic PD (*5*), (b) these pathogenic mutations increase kinase activity, which is required for neuronal toxicity (*6*) (*7*), (c) common variants in the *LRRK2* gene locus are risk factors for PD (*8*) (*9*), implicating a possible role in more common, currently idiopathic forms of PD, and (d) LRRK2 contains a kinase domain, which has been successfully manipulated through pharmacology. Concerns around potential safety liability of LRRK2-targeted therapies have emerged due to studies of rodent models of genetic deficiency in LRRK2, which show age-dependent emergence of phenotypes in kidneys and lungs (*10*) (*11*) *(*12*)* (*13*); Furthermore, repeated dosing of non-human primates with two LRRK2 kinase inhibitors (GNE-7915 and GNE-0877) induced microscopic, morphological changes in the lung similar to those observed in the rodent genetic models (*1*).

To thoroughly investigate the implications of possible lung effects on therapeutic development of LRRK2 kinase inhibitors, we designed a set of non-human primate studies to answer several questions (Figure 1). First, we asked whether the observed lung effects were attributable to on-target LRRK2 kinase inhibition or an off-target effect of the test compound. Since Fuji et al (2015) tested two structurally similar molecules, we designed a study to compare the effects of GNE-7915 with two additional LRRK2 kinase inhibitors (MLi-2, PFE-360) whose structures and off-target activities were distinct from each other as well as from GNE-7915. Second, we sought to determine if there was a safety margin for this lung effect using estimated clinically relevant exposures based on target inhibition in the brain. To address this, we tested MLi-2 and PFE-360 at two dose levels to determine whether a no-effect dose level for the lung finding can be observed concurrently with adequate brain LRRK2 kinase inhibition. Third, we asked whether the lung changes associated with LRRK2 kinase inhibition was reversible. To address this question, we included a dose-free recovery arm for GNE-7915 and MLi-2. Finally, we asked whether the observed lung phenotype has any measurable effect on pulmonary function by examining the effects of GNE-7915 and MLi-2 on pulmonary function in non-human primates using the same dosing regimens that were employed to assess lung toxicity.

**Figure 1:**
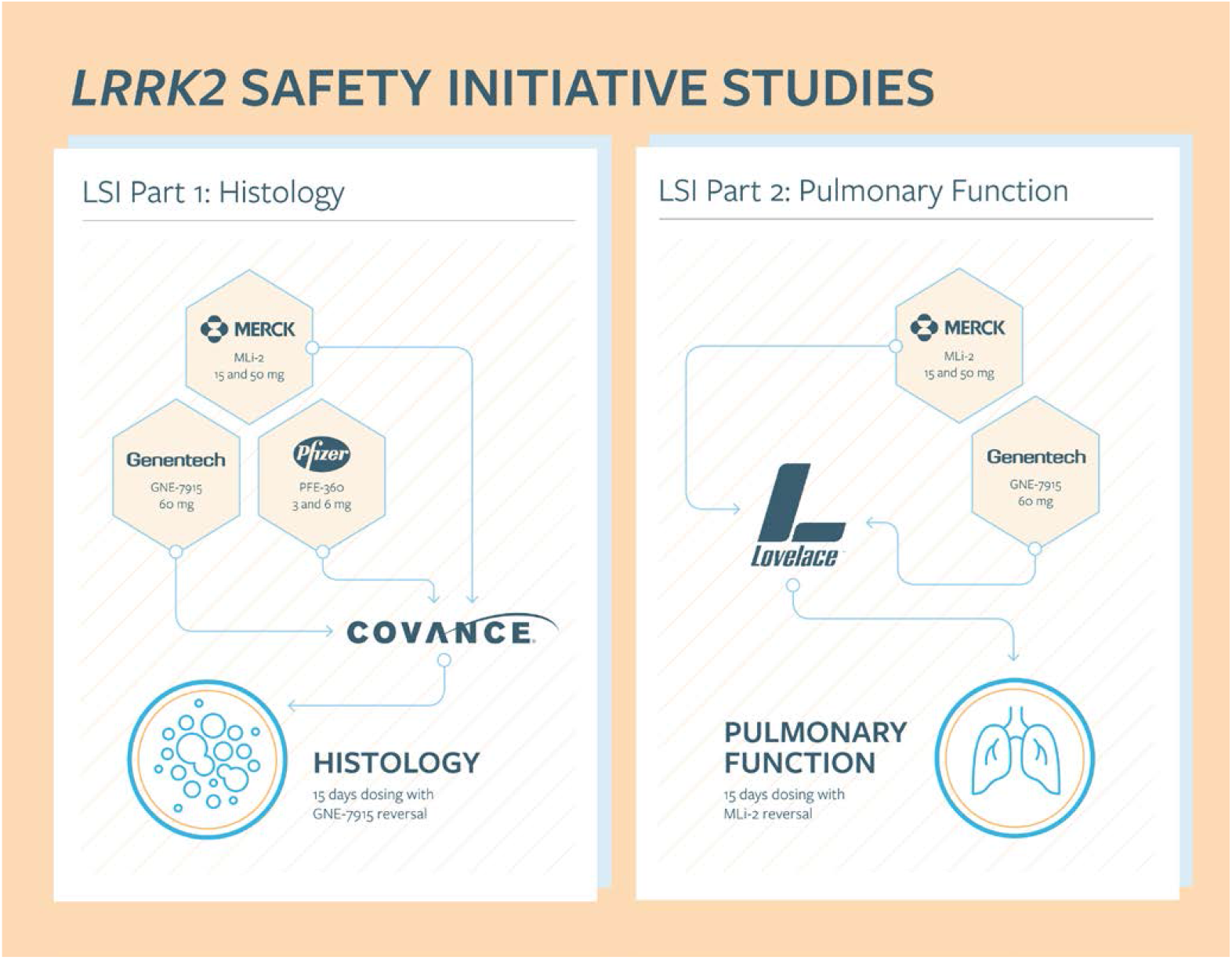
Schematic of the overall design of the LRRK2 Safety Initiative Studies

Through assessment of plasma and tissue exposures, pharmacodynamic effects, and light and electron microscopic evaluations of the lung, we demonstrate that two weeks of dosing with all three LRRK2 kinase inhibitors produce the same morphological changes in type II pneumocytes which were classified as minimal to mild in severity in prior reports (*1*). These data support that the changes observed in lung histology are due to LRRK2 kinase inhibition, however the lack of associated pulmonary functional impairment, reversibility, and the demonstration of a no effect dose with maintenance of CNS target engagement support continued investigation of LRRK2 kinase inhibition for PD.

## Results

### Part 1: Repeat-dose toxicological assessment

Doses for all LRRK2 inhibitors were chosen to match and expand on the exposure multiples achieved by GNE-7915 in previous work by Fuji et al. 2015. This was calculated by taking the (predicted monkey unbound plasma AUC over 24 hours) / (the unbound concentrations equal to IC50 for pS935-LRRK2 * 24). We aimed for a 1-10X range of exposure multiples over the compounds’ IC50 in mouse brain.

All compounds were well-tolerated and clinical observations in the GNE-7915-treated animals were similar to those previously reported (*1*). Lungs of animals administered GNE-7915 (30 mg/kg/dose), higher doses of MLi-2 (50 mg/kg, QD) and higher doses of PFE-360 (6 mg/kg, QD) showed slight increases in cytoplasmic vacuolation of type II pneumocytes, consistent with the effect previously seen with GNE-7915. None of the animals treated with lower doses of MLi-2 (15 mg/kg, QD) or PFE-360 (3 mg/kg, QD) showed the lung phenotype in this study. Two weeks after cessation of GNE-7915 dosing, type II pneumocyte vacuolation had returned to control levels, indicating complete reversibility (Figure 2). The present study established a no-effect dose level for both MLi-2 and PFE-360. Neither kidney nor brain showed any histopathological or macroscopic changes in response to any of the tested inhibitors, respectively.

**Figure 2:**
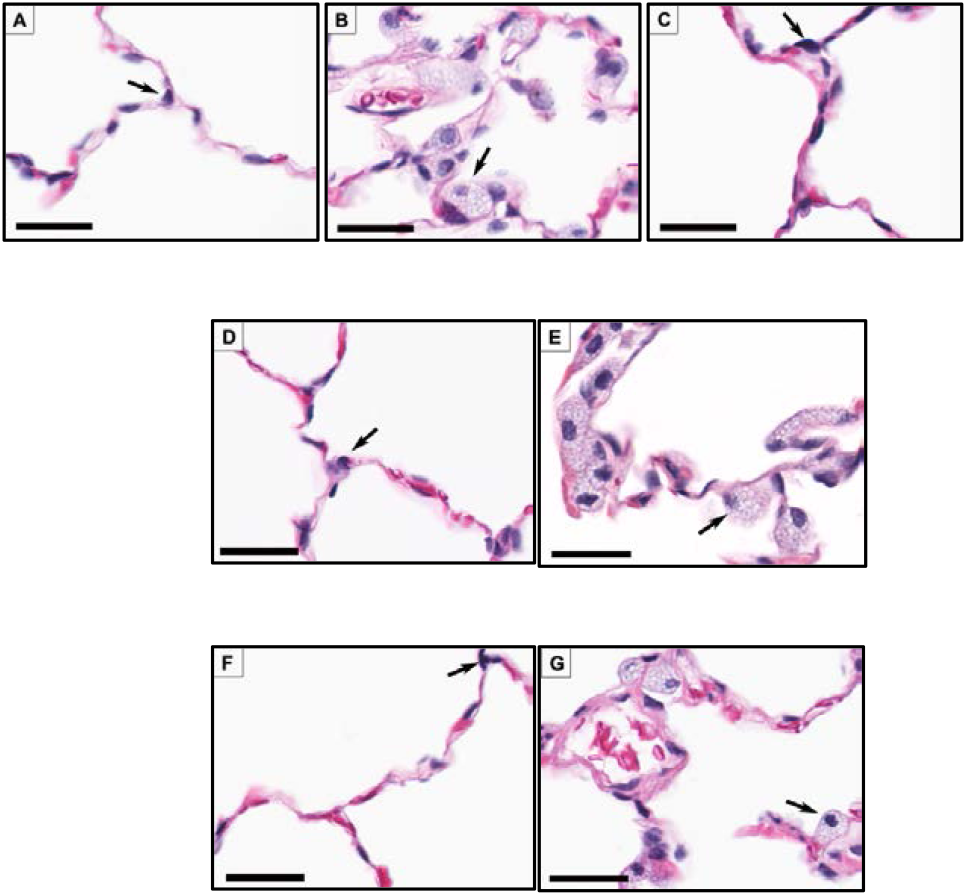
Effect of GNE-7915, MLi-2, and PFE-360 on type II pneumocytes to demonstrate a no effect dose level and reversal. Representative light microscopic images of hematoxylin and eosin stained sections of lung from vehicle control (A), GNE-7915 (B) GNE-7915 after 2-week recovery (C), low dose MLi-2 (D), high dose MLi-2 (E), low dose PFE-360 (F) and high dose PFE-360 (G). Note the increased cytoplasmic vacuolation of pneumocytes (arrows) in GNE-7915 (B) and high doses of MLi-2 (E) and PFE-360 (G) compared with vehicle control (A). Low dose MLi-2 (D) and PFE-360 (F) appear comparable to vehicle control (A) demonstrating a no effect dose level. Also note that GNE-7915 recovery group (C) is also comparable to vehicle control (A) demonstrating complete reversal of this change after a two-week drug free period. Bar = 30 mm.

To further characterize the lung changes, we examined tissues from a subset of animals (total five animals: one male from the vehicle control group, one male plus one female each from the high dose PFE-360 and high dose MLi-2 group) by Transmission Electron Microscopy (TEM). In the vehicle-treated animal, Type II pneumocytes exhibited typical ultrastructural appearance with the presence of cytoplasmic lamellar bodies characterized by membrane-bound vacuoles containing concentric laminations of electron-dense membranous material (*14*). Similar to the H&E staining, animals treated with MLi-2 at 50 mg/kg or PFE-360 at the 6 mg/kg showed a qualitative increase in the number and size of lamellar bodies (Figure 3). Despite this prominent morphological finding there were no associated ultrastructural findings indicative of degeneration. These ultrastructural data correlated with the light microscopic observation of increased pneumocyte vacuolation and confirmed that the affected cells were type II pneumocytes based on the presence of characteristic lamellar bodies.

**Figure 3:**
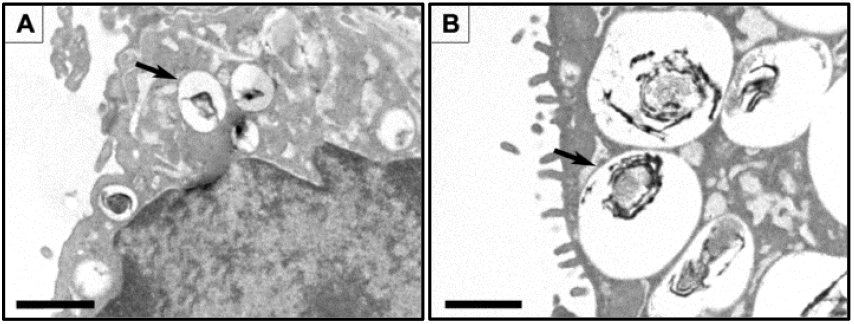
Representative ultrastructural images of type II pneumocytes demonstrating a qualitative increase in the size and number of lamellar bodies. Representative images of the lung section from a vehicle control (A) and a high dose MLi-2 (B) treated nonhuman primate are shown (note that PFE-360 animals showed similar qualitative changes). TEM demonstrate that the vacuolated cells noted by light microscopy are type II pneumocytes and that the test article-related increase in vacuolation is due to a qualitative increase in the size and number of cytoplasmic lamellar bodies (arrows), which are organelles normally present in this cell type. Bar = 1 mM

### Toxicokinetic analysis for plasma exposure multiples

Both MLi-2 and PFE-360 showed dose-related increases in plasma exposures (Table 1) For GNE-7915, the observed AUC for unbound plasma level was somewhat higher and resulted in 2-3.2x the exposure multiple over unbound IC_50_ for pSer935-LRRK2 reduction in the mouse brain. For the low and high dose of PFE-360, the observed AUCs were slightly lower or close to the targeted exposure multiple, respectively. Notably, the low-dose exposure multiples of 1-2x were in a range close to those seen in the GNE-7915 group. In contrast, the observed unbound plasma AUCs for MLi-2 were significantly higher than the 1x and 10x multiples targeted. Thus MLi-2 produced exposure multiples of 9.5x to 16.9x at the low dose and 45.5x to 64.8x at the high dose.

**Table 1:**
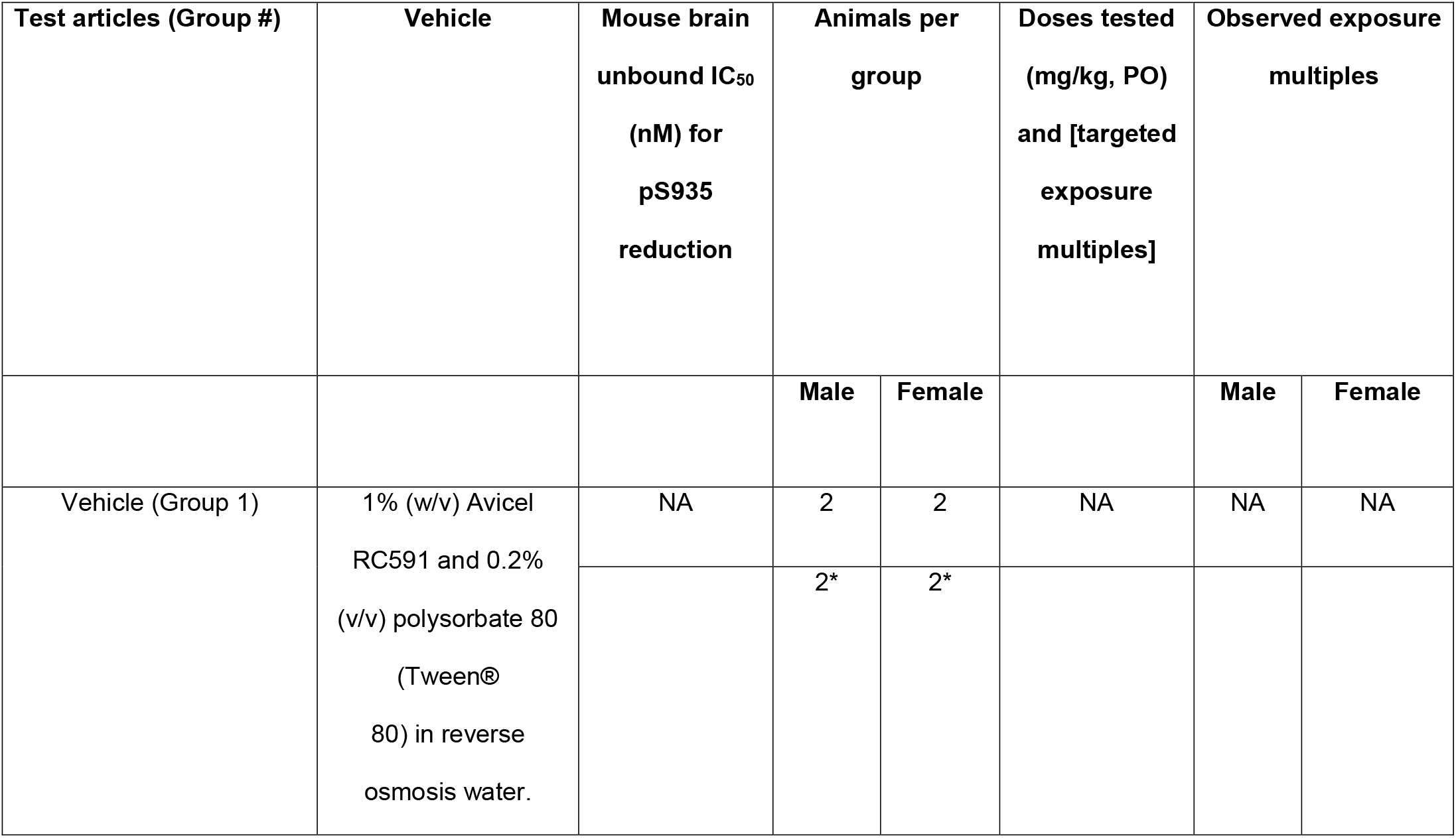
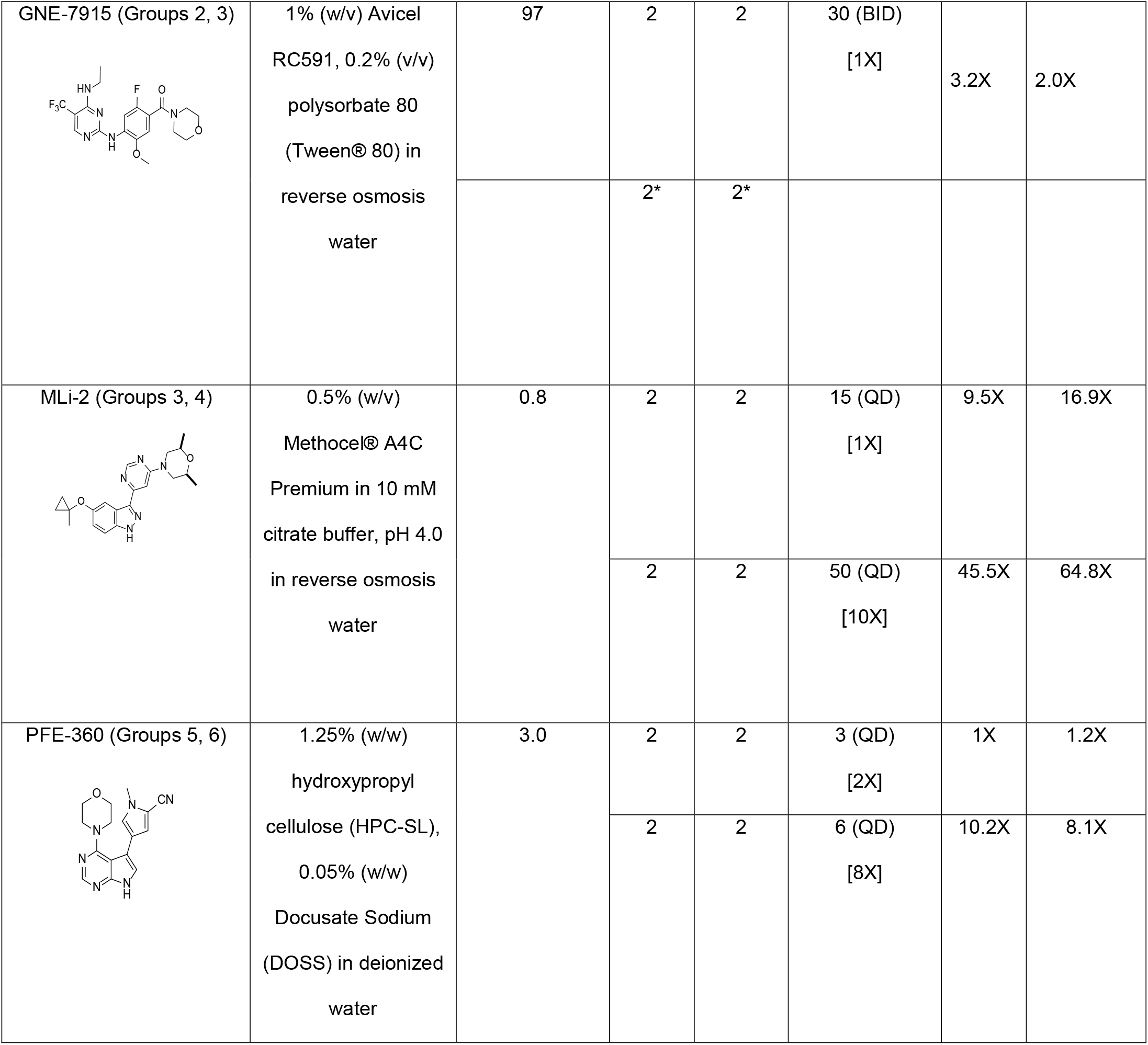
Dosing regimen and associated exposures of LRRK2 kinase inhibitors. Dosing regimens of each compound were selected based on PK/PD studies conducted at each company. Expected versus observed exposure multiples for each test compounds are shown. *Indicates animals used to assess recovery.

| Test articles (Group #) | Vehicle | Mouse brain<br>unbound IC<sub>50</sub><br>(nM) for<br>pS935<br>reduction | Animals per<br>group |  | Doses tested<br>(mg/kg, PO)<br>and [targeted<br>exposure<br>multiples] | Observed exposure<br>multiples |  |
| --- | --- | --- | --- | --- | --- | --- | --- |
|  |  |  | Male | Female |  | Male | Female |
| Vehicle (Group 1) | 1% (w/v) Avicel<br>RC591 and 0.2%<br>(v/v) polysorbate 80<br>(Tween®<br>80) in reverse<br>osmosis water. | NA | 2 | 2 | NA | NA | NA |
|  |  |  | 2* | 2* |  |  |  |
| **GNE-7915 (Groups 2, 3)** | 1% (w/v) Avicel<br>RC591, 0.2% (v/v)<br>polysorbate 80<br>(Tween® 80) in<br>reverse osmosis<br>water | 97 | 2 | 2 | 30 (BID)<br>[1X] | 3.2X | 2.0X |
|  |  |  | 2* | 2* |  |  |  |
| **MLi-2 (Groups 3, 4)** | 0.5% (w/v)<br>Methocel® A4C<br>Premium in 10 mM<br>citrate buffer, pH 4.0<br>in reverse osmosis<br>water | 0.8 | 2 | 2 | 15 (QD)<br>[1X] | 9.5X | 16.9X |
|  |  |  | 2 | 2 | 50 (QD)<br>[10X] | 45.5X | 64.8X |
| PFE-360 (Groups 5, 6) | 1.25% (w/w)<br>hydroxypropyl<br>cellulose (HPC-SL),<br>0.05% (w/w)<br>Docusate Sodium<br>(DOSS) in deionized<br>water | 3.0 | 2 | 2 | 3 (QD)<br>[2X] | 1X | 1.2X |
|  |  |  | 2 | 2 | 6 (QD)<br>[8X] | 10.2X | 8.1X |

**Table 2:** Total tissue concentrations of GNE-7915, MLi-2 and PFE-360 in NHPs following 15 day’s treatment expressed as a ratio of total plasma concentration.

| **Compound** | **Dose** | **[Brain]/[Plasma]** | **[Lung]/[plasma]** |
| --- | --- | --- | --- |
|  | **(mg/kg)** | **ratio** | **ratio** |
| **GNE-7915** | 30 | 3.7± 0.52 | 5.26±1.04 |
| **MLi-2** | 15 | 0.33±0.03 | 0.75±0.02 |
|  | 50 | 1.01±0.46 | 2.70±1.07 |
| **PFE-360** | 3 | 0.47±0.16 | 4.05±1.98 |
|  | 6 | 0.57±0.03 | 2.04±0.33 |

Supplemental table 2 shows total concentrations of GNE-7915, MLi-2 and PFE-360 in the brain and lung expressed as a ratio of the total plasma concentration. These data indicate that there were no differences in lung/plasma ratios between these compounds.

### Target inhibition (pSer935-LRRK2 and pT73-Rab10)

All three compounds tested demonstrated LRRK2 kinase activity in the lung, PBMCs and brain, as assessed by a reduction in the ratio of pSer935-LRRK2/total LRRK2 levels *ex vivo* (Figure 4). In the lung and PBMCs, LRRK2 kinase activity was completely inhibited by GNE-7915, MLi-2 and PFE-360 at all doses tested. In the brain, GNE-7915 and MLi-2 (15 and 50 mg/kg, QD) and PFE-360 (6 mg/kg, QD) reduced pS935-LRRK2/total LRRK2 but only a partial reduction was seen at the low dose of PFE-360 (3 mg/kg, QD). After a two-week withdrawal of GNE-7915, levels of pSer935-LRRK2 in the lung, PBMC and brain returned to levels comparable to the vehicle-treated group. Treatment with all three LRRK2 kinase inhibitors produced variable but statistically significant reductions in total LRRK2 levels in the brain and PBMCs but not the lungs.

**Figure 4:**
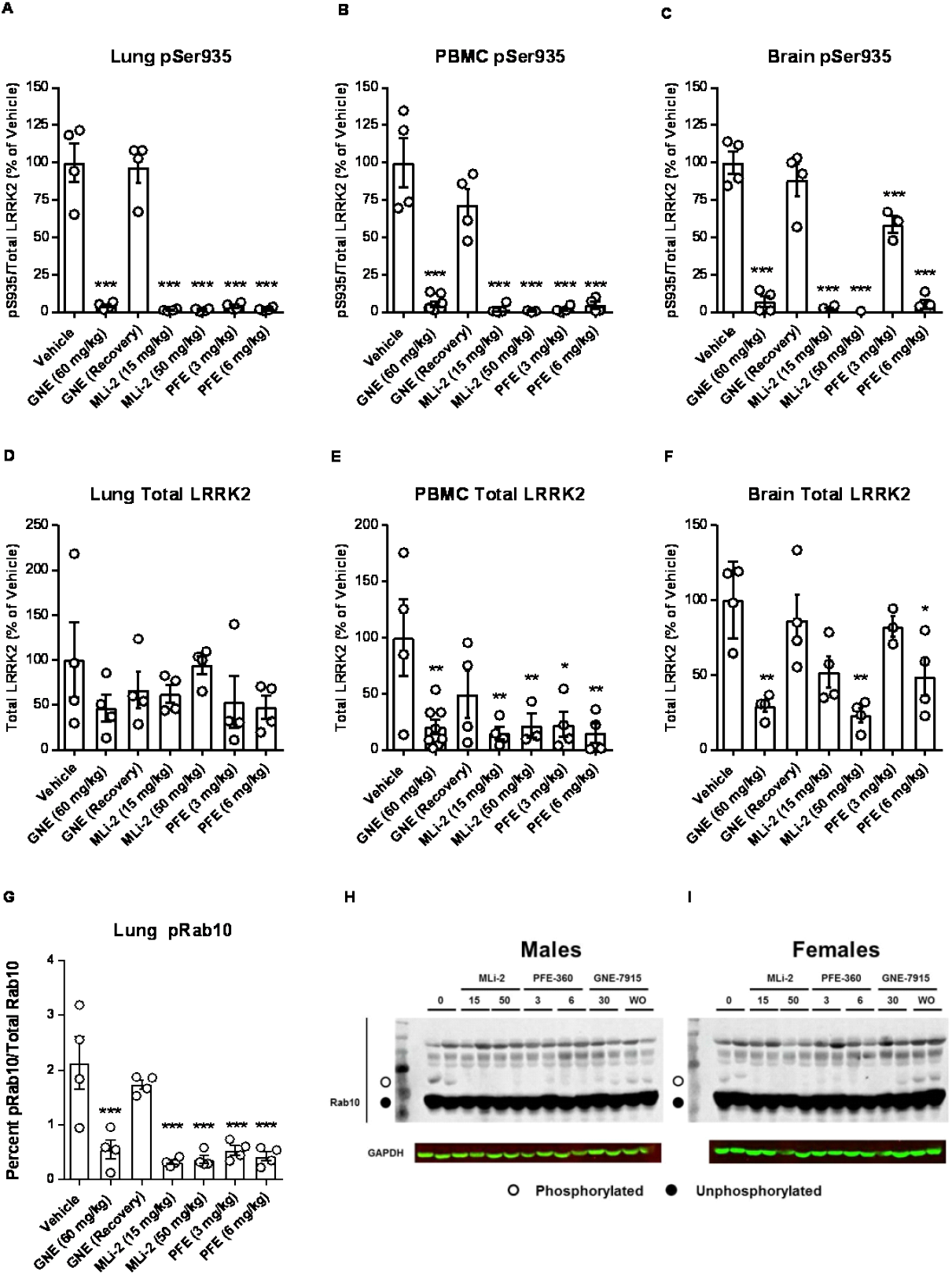
Target inhibition in lungs, PBMC and brain. Effects if LRRK2 kinase inhibitors on pSer935 and pRab10 levels were evaluated. pS935-LRRK2/total LRRK2 and total LRRK2/GAPDH ratios were measured to evaluate LRRK2 kinase inhibition in (A and D) lung (B and E) PBMC (C and F) after 15 days of dosing with GNE-7915 (30 mg/kg BID), MLi-2 (15 and 50 mg/kg QD) and PFE-360 (3 and 6 mg/kg QD). Percent pRab10/total Rab10 levels in the lung as determined by Phos-tag gel analysis is shown in G. pRab10/total Rab10 western blots for (H) males and (I) females. pSer935 and pRab10 data for animals from who GNE-7915 treatment was withdrawn (recovery arm) for 14 days is also shown. Data are mean ± SEM; n = 4-8 per treatment group. *, P < 0.05, **, P <0.01, ***, P < 0.001 compared with the vehicle treated animals.

Since pSer935 does not represent a direct phosphorylation site of LRRK2, we also evaluated the levels of pThr73 Rab10, identified as an *in vivo* substrate for LRRK2 kinase *(*15*)* and thereby offering a better indicator of LRRK2 kinase activity. All three LRRK2 inhibitors significantly reduced pThr73 Rab10 levels, assessed using the Phos-tag system, in the lung without affecting total Rab10 levels. Notably, both low and high doses MLi-2 and PFE-360 produced similar reductions in pThr73 Rab10. As with pSer935 LRRK2, we saw a complete recovery in pThr73 Rab10 levels to those seen in vehicle controls in the lungs of animals two-weeks after withdrawal of GNE-7915 (Figure 4).

### Relationship between the observed unbound plasma concentration and IC50 for mouse brain pSer935 LRRK2 inhibition

Neither the observed exposure multiple (Table 1) nor the target inhibition data (Figure 4) completely explain the observation that the lung effect was seen only in the GNE-7915 and high dose MLi-2 and PFE-360 groups. To investigate whether lung changes could be due to the duration of LRRK2 kinase inhibition during the dosing cycle, we examined the relationship between unbound plasma exposure-time profile to the unbound plasma IC50 for mouse brain pSer935 LRRK2 inhibition for each compound (Figure 5). These data show that the minimum unbound plasma concentration (Cmin) for GNE-7915 remains above its IC50 for brain target inhibition (pS935-LRRK2) at all times over the course of 24 hours. In contrast, regression analyses of TK data indicate that the time over IC50 for unbound plasma exposures for the low doses of MLi-2 (15 mg/kg, QD) and PFE-360 (3 mg/kg, QD) was 17 hours and 8.5 hours, respectively. Notably, animal #105235 in the high dose MLi-2 group did not show a lung effect and had plasma exposures that appear to fall below the IC50 of 0.8 nM at a time point less than18 hours. These data lead us to hypothesize that the lung effect is likely to be mediated by Cmin and its relationship to IC50 for brain target inhibition. Considering the shape of the concentration-time profile in the terminal phase, the 17-hour threshold is likely an over estimate of the actual time at which the free plasma concentration fell below the IC50.

**Figure 5:**
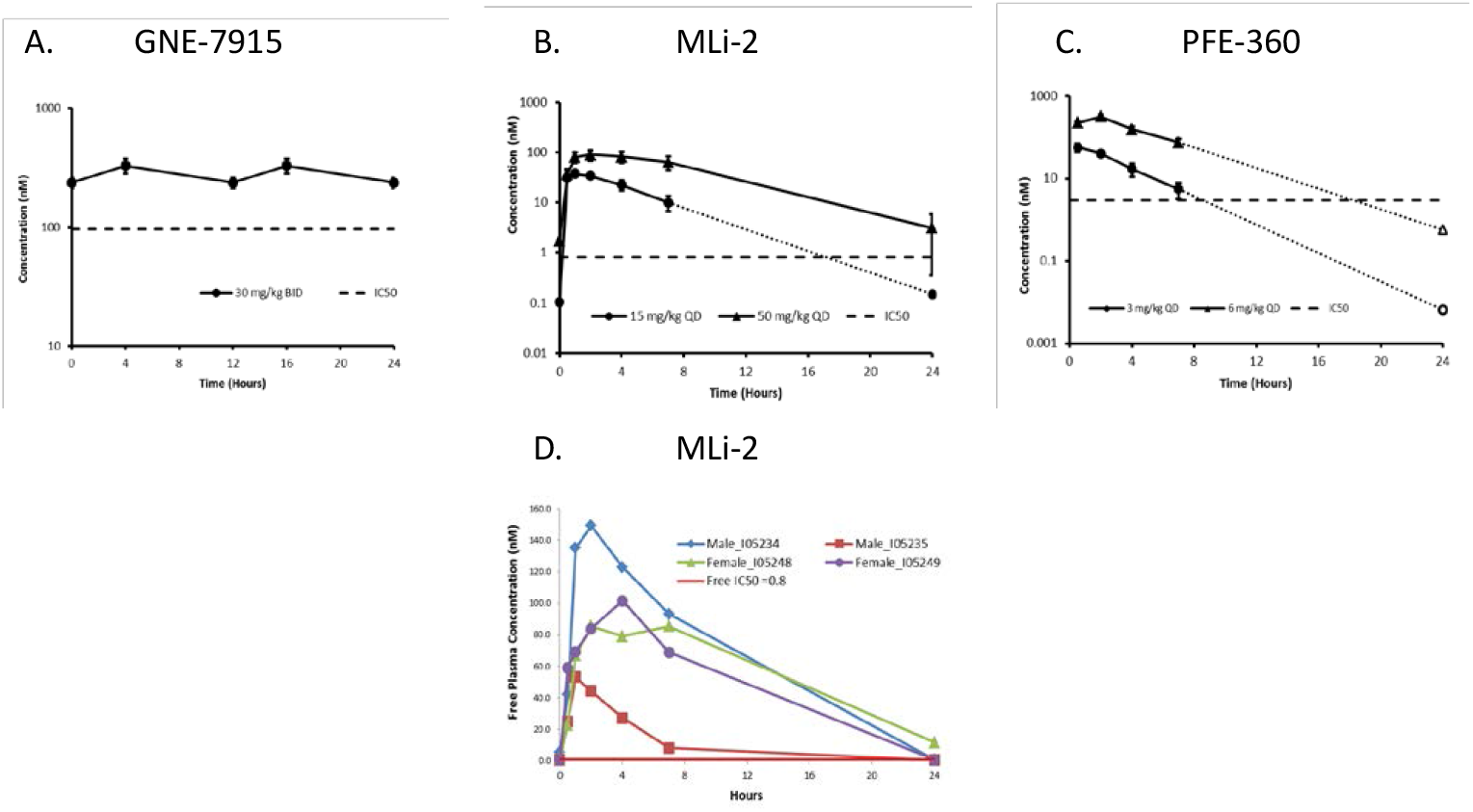
Relationship between the observed unbound plasma concentration and IC50 for mouse brain pS935-LRRK2 inhibition. The concentration-time profile (mean ± SEM) for unbound plasma levels for the GNE-7915 (panel A), MLi-2 (panel B) and PFE-360 (panel C) are shown. The closed symbols represent observed data while the dotted lines and open symbols represent extrapolated values using log-linear regression of the observed data from 2 to 7 hours. Regression analysis was applied since the majority of animals in these groups had plasma concentrations less than the lower limit of quantitation at 24 h. The dashed line in each panel represents the unbound IC50 for mouse brain pS935-LRRK2 inhibition for the respective compound. Panel D shows unbound plasma concentration-time profile for individual animals treated with 50 mg/kg MLi-2 to highlight animal #105235 (red line) who did not show the vacuolation of Type II pneumocytes in the lung while showing the shortest duration of time (18 h) during which the plasma exposures were above the IC50 for brain pS935-LRRK2 inhibition. The 18 hr threshold was estimated by log-linear regression between 7 and 24 hr. Considering the shape of the concentration-time profile in the terminal phase, 18 hr is likely an over estimate of the actual time on which the free plasma concentration fall below the IC50.

### Di-22:6-BMP levels in biofluids and tissues

Previous studies (*1*) showed that genetic deletion (LRRK2 knockout) and pharmacologic inhibition of LRRK2 kinase activity by GNE-7915 reduced urinary di-22:6-BMP levels. Consistent with the previous study, we observed significant decreases in di-22:6-BMP levels in the urine in all LRRK2 kinase inhibitor treated groups at the end of the dosing period compared to pre-dose samples (Figure 6). Following the two-week dose-free interval for animals administered GNE-7915, di-22:6-BMP levels returned to baseline ranges, indicating full reversibility. In contrast to the urine, di-22:6-BMP levels in brain, lung and kidney tissues were highly variable and failed to show a consistent pattern of changes induced by LRRK2 inhibitor treatment (data now shown). Although the source of urinary di-22:6-BMP remains unclear, the data shown here clearly demonstrate that this biomarker reflects on-target and reversible pharmacology of LRRK2 inhibitors. However, di-22:6 BMP does not differentiate with lung effects and therefore, is not suitable as a biomarker of the lung findings.

**Figure 6:**
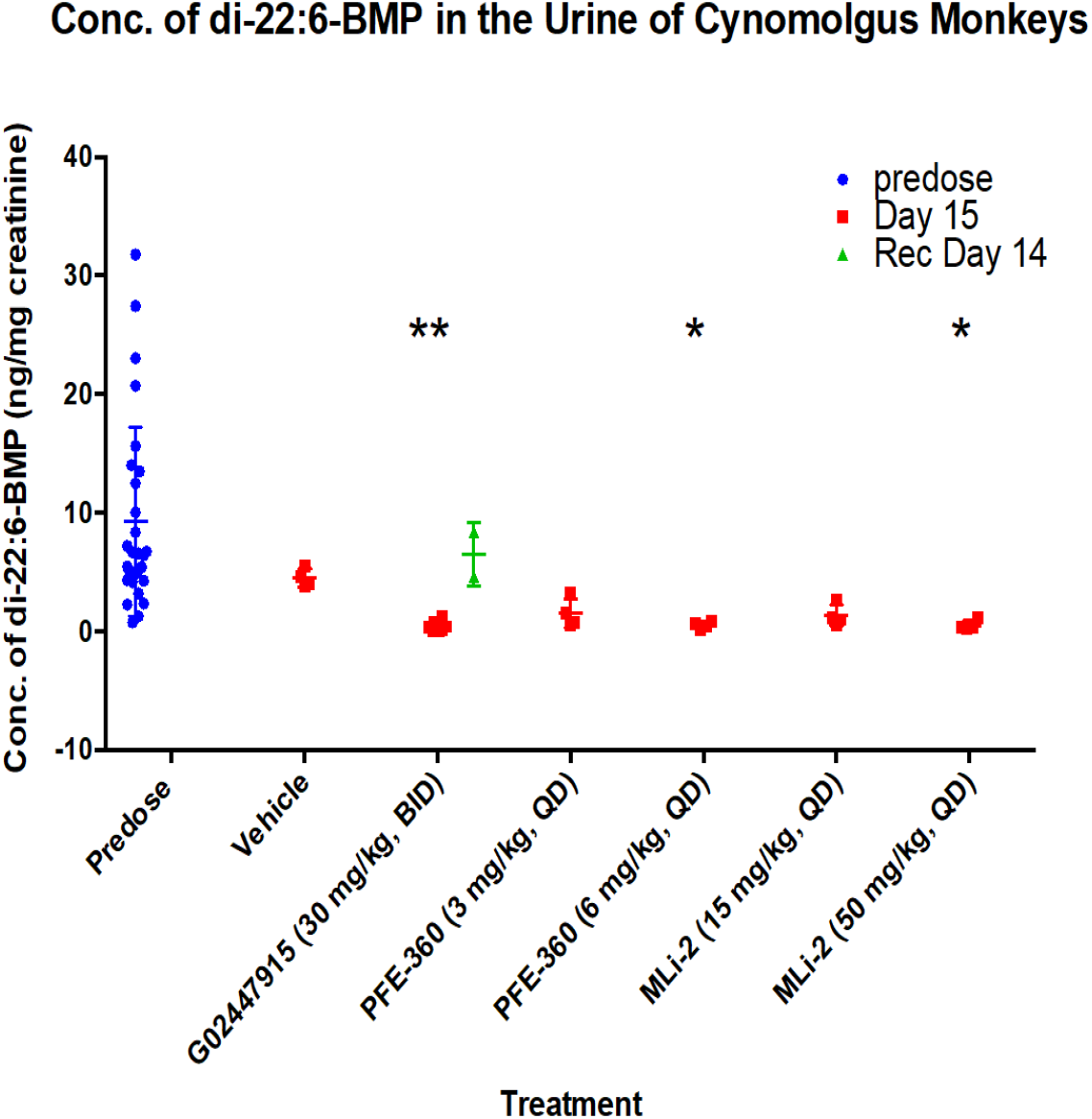
Concentration of di-22:6-BMP in the urine of cynomolgus monkeys. Levels of di-22:6 are shown for predose, and vehicle, GNE-7915 (30 mg/kg, BID), PFE-360 (3 mg/kg, QD and 6 mg/kg, QD), and MLi-2 (15 mg/kg, QD and 50 mg/kg, QD) on days 15 and recovery day 14. All LRRK2 kinase inhibitors significantly decreased urine di-22:6 BMP and returned to baseline following a two-week dose-free recovery interval.

### Part 2: Effect of MLi-2 and GNE-7915 on Lung Function

In a separate cohort of non-human primates, we administered high and low doses of MLi-2 (15 mg/kg and 50 mg/kg, QD) daily for 15 days as in our earlier studies and (as in Part 1) were evaluated effects on lung function during the dosing phase as well as after a two-week recovery phase (see schematic of the experimental design in Figure 10).

### Lung Histopathology

On day 15, all four animals treated with high doses of MLi-2 (50 mg/kg) showed a widespread increase in cytoplasmic vacuolation of type II pneumocytes similar to the changes observed in our earlier studies. Pathological examination graded these alterations as minimal in three animals and mild in one animal. In contrast, only one of four animals administered low doses MLi-2 (15 mg/kg) showed a wide-spread increase in cytoplasmic vacuolation of type II pneumocytes. Of the remaining three animals, two showed no changes in the lung whereas one animal had a few scattered type II pneumocytes with increased cytoplasmic vacuolation. Following the 13-day recovery period, histological examination of lungs of each animal in both the high and low dose groups were unremarkable, indicating complete recovery of the lung effect as was seen earlier with GNE-7915.

Minimal to mild vacuolation to type II pneumocytes was observed in all eight GNE-7915-dosed animals as expected.

### Pulmonary Functional Tests

Lung function tests were conducted at four different time points: baseline (day 0), day 7 and day 15 of dosing phase, and day 28 representing the two-week withdrawal from the treatment. Neither dose of MLi-2 affected lung diffusion capacity for carbon monoxide (DLCO) or elasticity of the lung, as determined by quasi-static lung compliance in the insufflation pressure range of 0 to 10 cmH2O (Cqs10) (Figure 7). Similarly, MLi-2 did not affect forced vital capacity (FVC), forced expiratory volumes (FEV) or forced expiratory flows (FEF) and mean mid-expiratory flows (MMEF) (Figure 7). MLi-2 also had no apparent effect on ventilator capacity as determined by forced expiratory flows given the lack of change over time and that ventilator capacity values in treated animals generally fell within the range of variability exhibited.

We also tested GNE-7915 (30 mg/kg, BID) at baseline, day 7, and day 15 of treatment. As with MLi-2, animals treated with GNE-7915 did not differ from the vehicle control group in any of the tests administered (data not shown).

**Figure 7:**
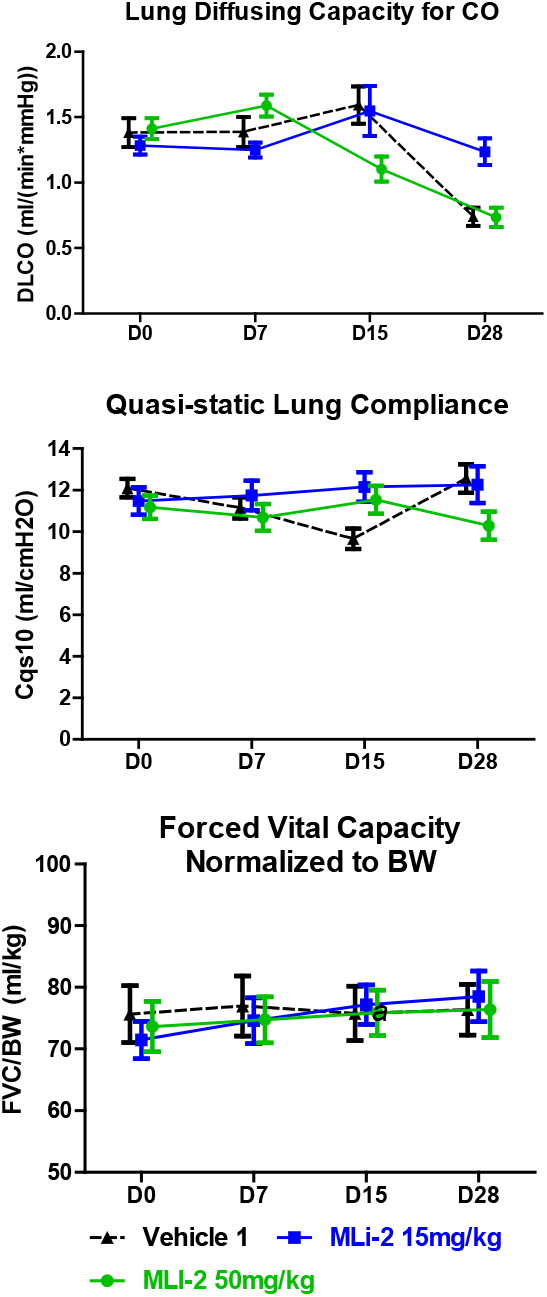
Pulmonary Function Tests for MLi-2. Lung diffusion capacity for carbon monoxide (DLCO), elasticity of the lung, as determined by quasi-static lung compliance in the insufflation pressure range of 0 to 10 cmH2O (Cqs10) and forced vital capacity (FVC) were not affected by both doses of MLi-2. GNE-7915 (30 mg/kg, BID) also had no effect on these functional tests (data not shown).

### Toxicokinetic data

For both MLi-2 and GNE-7915, exposure multiples were calculated using the observed unbound AUC, 0-8 hr and unbound IC50 for pSer935-LRRK2 * 8 hr. Consistent with the toxicokinetic data for MLi-2 in part 1, MLi-2 showed dose-related increases in plasma exposures and the unbound plasma AUCs were significantly higher than the 1x and 10x multiples targeted. On day 8 the unbound AUC for MLi-2 produced exposure multiples of 20x at the low dose and 35x at the high dose. For GNE-7915, the observed AUC for unbound plasma level was somewhat higher and resulted in 2x the exposure multiple over unbound IC50.

### Target inhibition (pThr73 Rab10)

Using a phospho-specific antibody, we assessed levels of pThr73 Rab10 in lung tissue from animals dosed with vehicle or GNE-7915 (30mg/kg) twice daily for 15 days. Levels of pThr73 Rab10 were significantly decreased in samples from GNE-7915 dosed animals compared to vehicle controls while total Rab10 levels were comparable between both groups (Figure 9). These data show significant inhibition of LRRK2 activity in animals dosed with GNE7915 and demonstrate that pThr73 Rab10 levels can be used as a pharmacodynamic readout of LRRK2 activity *in vivo*.

### Effect of MLi-2and GNE-7915 on Bronchoalveolar Lavage

Phosphatidylcholine levels after MLi-2 treatment did not differ from vehicle-treated animals suggesting that the compound did not alter surfactant release (Figure 8). MLi-2 had no effect on nucleated cell count suggesting no inflammatory response. Although some variability in phosphatidylcholine levels was noted at the low dose (15mg/kg) of MLi-2 this was not an effect of the test article given the absence of dose relationship (e.g., the high dose (50 mg/kg) did not differ from vehicle controls). Similarly, GNE-7915 did not affect surfactant release as assessed by phosphatidylcholine levels in the bronchoalveolar lavage fluid (data not shown).

**Figure 8:**
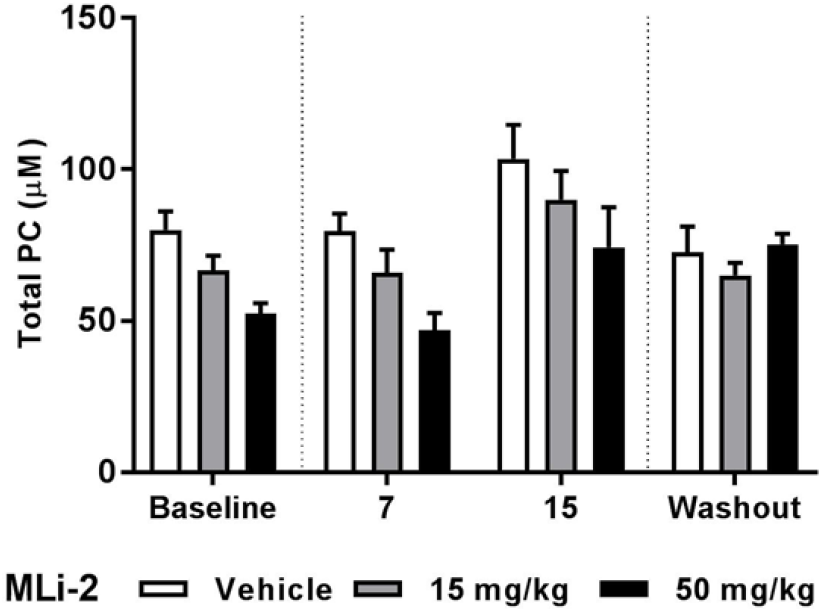
Effect of 15 days treatment with MLi-2 on phosphatidylcholine levels in BAL fluid from NHPs. BAL fluid was collected at baseline (day 0, prior to treatment initiation), day 7, day 15 and after compound withdrawal on day 28. Phosphatidylcholine levels in the BAL fluid at baseline (n=12 per group), day 7 (n=12 per group), 15 (n =12 per group) and 28 (n=8 per group) are presented as mean ± SEM.

**Figure 9:**
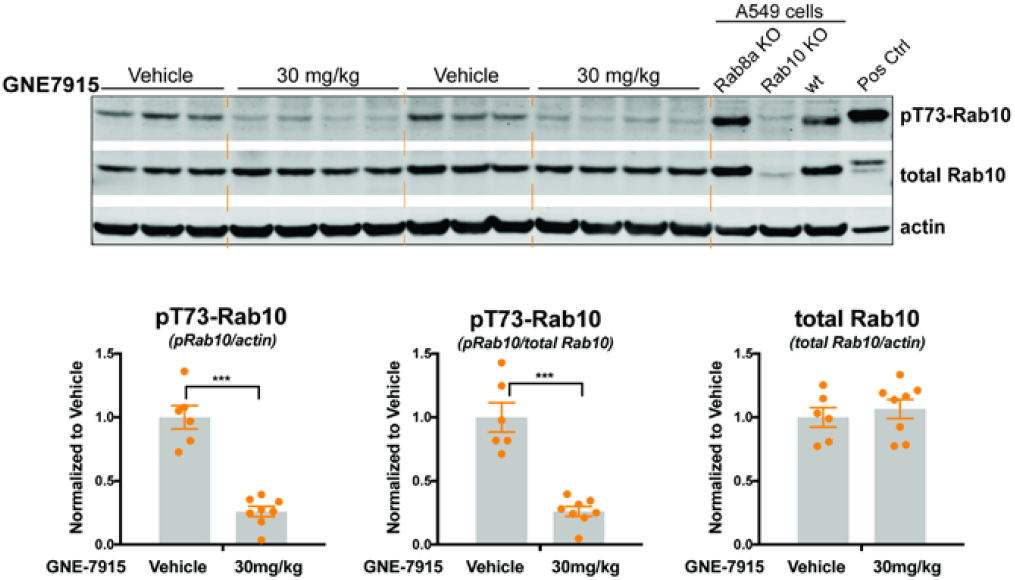
LRRK2 inhibitor, GNE-7915, significantly decreases pT73-Rab10 level in the lungs of cynomolgus monkeys. Using a phospho-specific antibody against pT73-Rab10, these results demonstrate that treatment with the LRRK2 inhibitor GNE7915 decreases pT73-Rab10 level in the lungs from in vivo dosed cynomolgus monkeys. A) Levels of pT73-Rab10 and total Rab10 in vehicle or GNE7915 dosed monkey lungs were assessed using western blot analysis. Levels of ß -actin were measured as a protein loading control. pRab10 levels were also assessed in wildtype, Rab8a KO, and Rab10 KO A549 cells to demonstrate the specificity of the pRab10 antibody. Positive control (shown as pos ctrl) lysates were generated from HEK293T cells co-expressing HA-Rab10 and FLAG-LRRK2-R1441C. B) pRab10, total Rab10, and ß -actin levels were quantified and normalized to signal seen in vehicle treated animals. pRab10 levels were decreased with GNE7915 treatment, while total Rab10 level were not affected; n=6 animals in the vehicle group and n=8 animals in GNE7915 dosed group. Error bars correspond to s.e.m. and p-value: t-test; *** p < 0.001.

## Discussion

The present studies demonstrate that repeated doses of three structurally distinct LRRK2 kinase inhibitors with different off-target pharmacologic profiles induces an identical response in the lungs of cynomolgus macaques. The lung effect is characterized by a minimal to mild increase in type II pneumocyte vacuolation that correlates ultrastructurally with an increase in size and number of cytoplasmic lamellar bodies in the absence of any associated degenerative findings. These data confirm previous studies (*1*) and support the conclusion that this effect is a consequence of LRRK2 kinase inhibition. However, several lines of additional evidence further clarify these results. First, kidney did not show any histopathological or macroscopic changes in response to any of the compounds. Second, we identified a clear no-effect dose of PFE-360 by demonstrating that PFE-360 inhibited LRRK2 kinase activity in the non-human primate brain at anticipated clinically relevant exposures without producing changes in the lung. For MLi-2, although we also observed a no-effect dose in one study, further studies showed that the low dose may be at the threshold of no-effect dose level since one out of twelve monkeys in our subsequent study showed lung vacuolation. These data indicate that it is possible to obtain a safety margin for this LRRK2 kinase inhibitor induced lung effect, an important consideration for ongoing and future therapeutic trials. Third, and importantly, a two-week withdrawal from treatment with either GNE-7915 or MLi-2 resulted in complete recovery of the type II pneumocyte vacuolation effect. Finally, a wide battery of clinically relevant pulmonary functional tests in non-human primates showed that two distinct compounds (MLi-2 and GNE-7915) at doses that produce the vacuolation microscopic changes in non-human primate lungs had no impact on pulmonary function nor did they affect surfactant secretion. Taken together, these data provide additional characterization of the potential mechanism-based liability of LRRK2 kinase inhibitors and help guide the preclinical and clinical development path for this class of compounds.

PFE-360, MLi-2 and GNE-7915 are chemically distinct LRRK2 kinase inhibitors that have minimal overlap in their off-target activity *(*16*) (*17*) (*18*) (*19*)* (Galatsis et al., in prep). Thus, a comparison of these three compounds offered us an opportunity to assess whether the lung phenotype might be mediated *via* LRRK2 kinase inhibition or other compound-specific off-target pharmacology. The *in vivo* plasma IC50 concentrations for inhibition of phosphorylation of pSer935 LRRK2 in wild-type mouse brain corrected for protein binding, are 98, 0.8 and 3.0 nM, respectively, for GNE-7915, MLi-2 and PFE-360. Although indirect, it is well established that pSer935 phosphorylation reflects LRRK2 kinase activity (*17*). Hence, we selected doses and a dosing regimen for MLi-2 and PFE-360 that were projected by non-human primate pharmacokinetic studies to achieve exposures that were either 1x-2x (low dose) or 8x-10x (high dose) the *in vivo* IC50 for pSer935 LRRK2 reduction in the mouse brain. GNE-7915 was used as an experimental positive control for this study, and therefore, its dosing was identical to that previously reported in Fuji et al., 2015 (*1*). Toxicokinetic analyses demonstrated that the observed exposures of PFE-360 and GNE-7915 were close to the targeted levels whereas those of MLi-2 were significantly higher. To directly assess whether the exposures achieved were sufficient for LRRK2 kinase inhibition in the brain, pSer935 LRRK2 and total LRRK2 levels were examined by immunoblots on striatal lysates. All compounds reduced pS935-LRRK2 in the striatum by over 90% at both doses tested. Small and somewhat variable reduction in total LRRK2 levels were also observed at all doses. Similarly, the lung and PBMCs also showed >90% inhibition of LRRK2 kinase activity in all treatment groups. Thus, the selected dosing paradigm resulted in tissue exposures sufficient for LRRK2 kinase inhibition at all doses.

Histopathology of lung tissue from both studies conducted here demonstrated the replication of the previously observed mild effect in the form of accumulation of lamellar bodies in type II pneumocytes in monkeys treated with 30 mg/kg of GNE-7915. Identical microscopic changes in the type II pneumocytes were observed in non-human primates treated with the high doses of PFE-360 (6 mg/kg, QD) and MLi-2 (50 mg/kg, QD). These data strongly indicate that the observed lung effects are attributable to the on-target (i.e., LRRK2 kinase inhibition) pharmacology of the three compounds. In contrast to the high dose effect, lower doses of MLi-2 and PFE-360, at exposures that significantly inhibited LRRK2 kinase activity in the non-human primate brain, produced minimal to no effect in the lungs. These data are consistent with previous findings (*1*) demonstrating that a lower dose of GNE-7915 did not produce the lung phenotype despite inhibiting LRRK2 activity in the brain of the non-human primates. Thus, it appears that a safety margin for this lung effect would be achievable with brain-penetrant LRRK2 kinase inhibitors.

The observed dose-dependent effects of MLi-2 and PFE-360 in the lung histology concurrently with >90% reduction in pSer935 LRRK2 levels in the lung by both doses of these compounds raised the question whether pSer935 LRRK2 is an adequate marker of LRRK2 kinase inhibition. This concern is especially critical since Ser935 phosphorylation is not induced directly by LRRK2 *(*20*)*. To circumvent this issue, we assessed the levels of a recently identified LRRK2 kinase target (*15*), pThr73 in Rab10 in the lung tissue (brain tissue was not examined since the available antibody is not able to detect brain pThr73 Rab10 levels at sufficient sensitivity and specificity). Interestingly, both low and high doses of MLi-2 and PFE-360 produced similar reductions in pThr73 Rab10 levels in the lungs without affecting total Rab10 levels. These data indicate that the target inhibition at a single time point cannot explain the observed sparing of the lungs at low doses of MLi-2 and PFE-360. Hence, we investigated whether the duration over which the compound exposures are maintained above their individual *in vivo* IC50 for brain pS935-LRRK2 may correlate with the observed lung changes. This analysis showed that while the levels of GNE-7915 remained above its IC50 for brain target inhibition over 24 h, the time over IC50 for the low doses of MLi-2 and PFE-360 were significantly shorter at 17 h and 8.5 h, respectively. The argument for time over IC50 was further bolstered by the observation that a single animal in the high dose MLi-2 achieved lower exposure and also failed to show lung changes. Thus, it appears that designing a dosing regimen that produces partial inhibition of LRRK2 in the lungs may be a viable strategy to avoid the induction of the cytoplasmic vacuolation in the type II pneumocytes. Of course, it is imperative to demonstrate that this strategy could produce the desired target inhibition and efficacy predicting end-points in the brain. It is also noteworthy that pathogenic mutations of LRRK2 increase kinase activity and emerging data indicate that LRRK2 kinase activity may be higher in idiopathic PD (*21*) (*22*). Thus restoration of LRRK2 kinase activity to physiological levels rather than more robustly inhibiting its activity may provide a viable therapeutic strategy with the potential of a better margin of safety.

In addition to the possibility of attaining a margin of safety for the lung effect via doses and dosing regimen, we demonstrate here, for the first time, that a relatively short (13-14 days) recovery phase results in complete amelioration of the lamellar body accumulation induced by two different compounds (GNE-7915 and MLi-2). In future studies, it will be important to study the time-course of recovery of the lung changes upon withdrawal from treatment with LRRK2 kinase inhibitors. It is also noteworthy that kidney tissues were free of any pathological changes, as was reported by Fuji et al. (2015).

We further explored the use of the FDA-approved biomarker, di-22:6-BMP, for cationic amphiphilic drug (CAD)-induced phospholipidosis as a safety biomarker for the lung change. Di-22:6-BMP is a phospholipid localized in the internal membrane of lysosomes and late endosomes where it is thought to participate in lysosomal degradation pathway (*23*). While no compound is a CAD, the hypothesis was that given the similarity of the LRRK2 inhibition-induced lamellar body change to that induced by CADs, increases in di-22:6-BMP may be useful as a means of monitoring for this change. In contrast to effects of CADs, there were no changes in di-22:6-BMP levels in the plasma while the urine showed a decrease in di-22:6-BMP levels. As such, the changes in urinary di-22:6-BMP levels appear to reflect a LRRK2-driven effect on lysosomal biology but is not likely to be useful as a safety biomarker. The di-22:6-BMP levels in the GNE-7915 group given a 2-week dose-free recovery period are within the range of baseline and/or vehicle control animals providing further evidence that the LRRK2 lysosomal effect was clearly reversible.

A major objective of our study was to go beyond the histopathological analyses and investigate functional effects of the observed microscopic changes in the lung. Using a comprehensive battery of pulmonary functional tests, we have established that LRRK2 kinase inhibition by GNE-7915 or MLi-2 at doses that induce an increase in size and/or number of lamellar bodies in type II pneumocytes does not disrupt pulmonary function. It is important to point out that the tests used here have a clinical correlate and thereby afford the ability to monitor lung function in clinical studies of LRRK2 inhibitors, should that be deemed necessary.

In summary, the studies described here have demonstrated that LRRK2 kinase inhibition can produce a minimal to mild, non-degenerative and reversible lung change in non-human primates after two weeks of treatment. These changes in the lung are not associated with any functional consequences and a safety margin for this effect appears to be achievable given the demonstration of no effect doses even with significant LRRK2 inhibition in the brain. Overall, these data suggest that this on-target morphological change in the lung of non-human primates treated with LRRK2 kinase inhibitor does not preclude advancing LRRK2 kinase inhibitors to the next stage of therapeutic drug development, including clinical evaluation in PD patients.

## Methods

### Animal welfare

For all animal studies reported here all procedures performed on animals were in accordance with regulations and established guidelines and were reviewed and approved by an Institutional Animal Care and Use Committee.

Part I study was conducted at Covance Laboratories, Inc., in Madison, WI. Monkeys had free access to water and were fed Certified Primate Diet #2055C (Harlan Laboratories, Inc.) one or two times daily and given various cage-enrichment devices and fruits, vegetables, or dietary enrichment for the duration of the studies.

### Part 1: Repeat-dose toxicological assessment

#### Animals and study design

The in-life phase, necropsy, biospecimens collection, anatomic pathology evaluation, and PK assessment were conducted at Covance Laboratories, Inc., Madison, WI. Studies were conducted in 28 Asian cynomolgus monkeys *(Macaca fascicularis)* aged 3 to 4 years and weighing 2 to 5 kg at the start of the study (Covance Research Products, Inc). The schedule of dosing, monitoring, necropsies and tissue collections are detailed in Table 1 and Supplement 1.

#### Doses and dosing regimen

The dose and dosing schedules were based on PK-PD studies and associated modeling of each compound conducted by participating companies. The in vivo potency of each molecule was determined by assessment of male C57Bl/6 mice (weighing 20-25g, 8-10 weeks old) brain pS935 reduction and expressed as unbound IC50s. The IC50 values were then used to calculate exposure multiples in monkeys by the formula: (predicted monkey unbound plasma AUC over 24 hours) / (the unbound concentrations equal to IC50 for pS935-LRRK2 * 24). The low doses of MLi-2 and PFE-360 were predicted to produce plasma exposure multiple of at least 1X, similar to that of GNE-7915, and the high doses were targeted to reach exposure multiples of 8-10X (Table 1).

#### Biospecimen Collection for Analyses of Compound Levels, In Situ Target Inhibition and Lipid Biomarker

Following sample collection for histopathology, additional biospecimens were collected as detailed below and in Table 1, weighed, snap-frozen in liquid nitrogen and placed on dry ice and stored at -60° to -80°C until processing for and various analyses.

##### Blood

Blood was collected from the femoral artery in separate aliquots: (a) in EDTA tubes to harvest plasma for analysis of toxicokinetics and the lipid biomarker (di-docosahexaenoyl (22:6) bis(monoacylglycerol) phosphate (di-22:6-BMP)) or (b) to isolate Peripheral Blood Mononuclear Cells (PBMCs) by density gradient separation using Ficoll-Paque PLUS (Sigma-Aldrich) for target inhibition assessment.

##### Cerebrospinal fluid (CSF)

CSF was collected from the cisterna magna at necropsy at approximately 4 h after the dosing to analyze compound levels.

##### Urine

Urinary aliquots were collected at baseline and approximately 4 h after dosing Day 15 and Day 28 (recovery phase) for analysis of di-22:6-BMP levels.

##### Brain, Lungs, Kidneys

At necropsy, brain, lungs and kidneys were collected unilaterally for compound exposures, target inhibition assessment and 22:6-BMP levels.

#### Compound levels in plasma, CSF and brain tissue

Plasma, CSF and tissue levels of compounds were determined by liquid chromatography and tandem mass spectrometry analyses (LC/MS/MS). The plasma and brain protein binding of test compounds were determined by equilibrium dialysis. All data are expressed as unbound compound level in each matrix.

#### Analyses of di-docosahexaenoyl (22:6) bis(monoacylglycerol) phosphate (di-22:6-BMP)

Plasma, urine, lung, kidney, and brain tissues from animals treated with all LRRK2 kinase inhibitors were analyzed for di-22:6-BMP levels by LC-MS/MS at Nextcea, Inc. (Woburn, MA) using extraction and analysis methods described in *(*24*)* (*25*).

#### Histopathology

Unilateral lung tissues (right median and diaphragmatic lobes) were collected at necropsy and immersion fixed in 10% neutral-buffered formalin (NBF) for at least 2, but no more than 3 days fixation before being transferred to 70% ethyl alcohol. Tissues were then processed to paraffin block and sectioned, at 4-6 um for routine hematoxylin and eosin staining and examination by light microscopy. Additional samples of lung were also collected in Karnovsky’s fixative and processed for evaluation by transmission electron microscopy as described previously (*1*).

#### Target inhibition assessment (pSer935-LRRK2 and pT73-Rab10 levels) by Western Blots

##### pS935-LRRK2 and total LRRK2

PBMC pellets and all other tissues were homogenized in lysis buffer (MSD Lysis Buffer) supplemented with protease (Roche cOmplete Mini) and phosphatase inhibitors tablets (Roche PhosStopTM). Lysates were centrifuged at 13.2 x 1000 rpm for 20 min at 4°C. Supernatants were collected and protein content per sample was determined by BCA colorimetric assay, using BSA as a standard (Life Technologies). 100 μg of protein was reduced (10% NuPage Reducing Agent and 5ul LDS Sample Buffer) and denatured at 70°C for 10 min, then resolved on 3-8% Tris-acetate gels (Life Technologies) and transferred to PVDF membranes (Life Technologies). Membranes were blocked in 5% dry milk in TBS plus Tween-20 (Sigma) for 1 hour at 4°C and probed with rabbit anti-LRRK2-pSer935 LRRK2 (Abcam; 1 μg/mL) overnight at 4°C. Membranes were then incubated with donkey anti-rabbit conjugated to HRP (Life Technologies; 1 μg/mL) combined with the IRDye 680CW (LI-COR, 926-68076) for 30 min at 4°C. LRRK2-pSer935-HRP signals were subsequently developed by luminol-enhanced chemiluminescence (SuperSignal^TM^ Substrate) and then visualized and analyzed on a LI-COR Odyssey system. For total LRRK2 detection, membranes were subsequently stripped (Life Technologies, Restore PLUS, 46430), re-blocked as above and probed with rabbit anti-LRRK2 antibody (Abcam, MJFF2 clone 41-2, AB133474; 1:500v/v) combined with mouse anti-GAPDH (Millipore, MAB374) overnight at 4°C. Membranes were then incubated with donkey anti-rabbit conjugated to HRP (Life Technologies, A16035; 1μg/mL) combined with the IRDye 800CW goat anti-mouse antibody (LI-COR) for 30 min at 4°C. HRP was then developed, visualized and analyzed as above. For quantification, pSer935-LRRK2 signals were normalized to total LRRK2. GAPDH levels were used for further normalization of protein loading for all samples to enable total LRRK2 quantification. pT73-Rab 10

#### Detection using Phos-tag gels

Lung tissue (500 mg) was homogenized in 500 □l of cold lysis buffer (50 mM Tris-HCl pH 7.5, 1%(v/v) Triton X-100, 50 mM NaF, 5 mM MgCl2, 10 mM β-glycerophosphate, 5 mM sodium pyrophosphate, 270 mM Sucrose,1mM sodium orthovanadate, 0.1 μg/ml mycrocystin-LR and Complete EDTA-free protease inhibitor cocktail). Lysates were centrifuged at 13.2 x 1000 rpm for 20 min at 4°C. Supernatants were collected, and protein concentrations were determined by Bradford assay using BSA as standard. Tissue lysates were denatured in Laemmli’s SDS-PAGE sample buffer (250 mM Tris-HCL PH6.8, 8% SDS w/v, 40% Glycerol v/v, 0.02% Bromophenol blue v/v and 4% 2-mercaptoethanol) at 70 °C for 10 minutes and spun at 13.2 x 1000 rpm for 1 minute at room temperature. 45 μg of lysates were loaded on to phostag gels (50 uM phostag, Supersep phos-tag ^TM^ 15% and 17 well, Wako Cat#196-16701) with WIDE-VIEW^TM^ Prestained Protein Size Marker III, run at 70V (40 minutes) for stacking and at 140V (3 Hours) in running buffer (25 mM Tris, 192 mM Glycine, 0.1% SDS, Wako Cat# 184-01291). Following washing, proteins were wet-transferred to nitrocellulose membranes. Membranes were blocked with 5% dry and skim milk dissolved in TBS containing 0.1% Tween. Membranes were probed with the primary, anti-Rab10 antibody (Cell Signaling, diluted 1:1000 with Solution One from SignalBoost™ Immunoreaction Enhancer Kit (EMD Millipore) containing 5% milk). Secondary antibody incubation was with anti-rabbit HRP-labeled IgG diluted 1:4000 in Solution 2 for one hour. GAPDH level was determined using mouse mAb (ab8245) and Li-Cor fluorescent 2^nd^ antibody. Images were developed for 3 minutes using SuperSignal West Dura Extended Duration substrate and captured by Odyssey Imager. The band intensity was measured and analyzed with image studio software. The total Rab10 level was defined by adding top pT73-Rab10 band and bottom Rab10 band intensity measurement. Percentage of phospho-Rab10 was calculated by (Phospho Rab/Total Rab) *100.

#### Detection using pT73-Rab10 specific antibody

Frozen non-human primate lung tissues (right cranial (RCr) region, 150-250 mg) was placed in pre-chilled 1.5 ml Eppendorf tube, with one Tungsten Carbide Beads (3 mm, Qiagen) and lysis buffer (5X of the tissue weight, e.g. 1000 ul lysis buffer for 200 mg tissue). The lysis buffer was composed of 50 mM Tris-HCl, pH7.4, 10 mM β-glycerophosphate, 10 mM sodium pyrophosphate, 0.1 μM mycrocystin-LR, 150 mM NaCl, 5 mM MgCl2, 1 mM EDTA, 1 mM DTT, 1% (v/v) Trion X-100, 10% glycerol, 100 nM GTPγS and cOmplete^TM^™ EDTA-free Protease Inhibitor Cocktail (Roche). The tissues were then homogenized for 6 minutes at 4 °C on TissueLyser II (Qiagen) at the frequency of 30/second, 3 min per time. Lysates were centrifuged at 14,000 rpm for 30 min at 4 °C. Supernatants were collected and protein concentrations were measured using Pierce™ BCA Protein Assay (ThermoFisher). Protein concentrations were normalized for equal protein loading and mixed with NuPAGE LDS Sample Buffer (4X, ThermoFisher) and NuPAGE Sample Reducing Agent (10X, ThermoFisher). Samples were incubated at 10 min at 70°C to denature proteins. 50 μg of tissue lysates were loaded onto NuPAGE™ 4-12% Bis-Tris Midi Protein Gels (ThermoFisher) and transferred to nitrocellulose membranes for 7 min using Trans-Blot^®^ Turbo™ Transfer

System (Bio-Rad). Membranes were blocked with Odyssey^®^ Blocking Buffer (TBS, LI-COR), incubated with primary antibody overnight at 4°C, and then incubated with secondary antibodies (1:20,000, LI-COR) for 1 hr at room temperature. The primary antibodies used were rabbit anti-pT73-Rab10 (1 μg/ml, polyclonal antibody, MJF-20, E8263, provided by Dr. Alessi), mouse anti-Rab10 (1 ug/ml, Acam, ab104859) and mouse anti-β-actin (1:5000, Sigma, A2228). LI-COR Odyssey system was used for western blot detection and quantitation. Western blot images were quantified using Image Studio software (LI-COR). The pT73-Rab10 or total Rab10 signal was normalized using the β-actin signal for each sample to control for loading. Data were plotted using GraphPad Prism 7.

#### Data analysis

Due to the small number of animals in this toxicological study, all toxicokinetic and histopathology data analyses were limited to the calculation of means and standard deviations without hypothesis testing via statistical methods such as regression or group comparisons.

The effect of the LRRK2 kinase inhibitors on pSer935 LRRK2 and pRab10 levels in the brain and peripheral tissues were analyzed by a one-way ANOVA followed by Dunnett’s multiple comparison test.

For di-22:6-BMP levels, unpaired t-tests were conducted to compare baseline measurements with the end of the dosing or recovery periods (plasma and urine), and to compare LRRK2 kinase inhibitor-dosed groups with the vehicle control group.

### Part 2: Lung Function Studies

A non-human primate study to evaluate pulmonary function was conducted at the Lovelace Respiratory Research Institute, Albuquerque, NM. The overall study design is shown in Figure 10. Full toxicokinetics (TK) analysis was not performed in this study in order to not disturb functional endpoints. Sparse TK performed in this study showed data consistent with the TK data in Part I.

**Figure 10.**
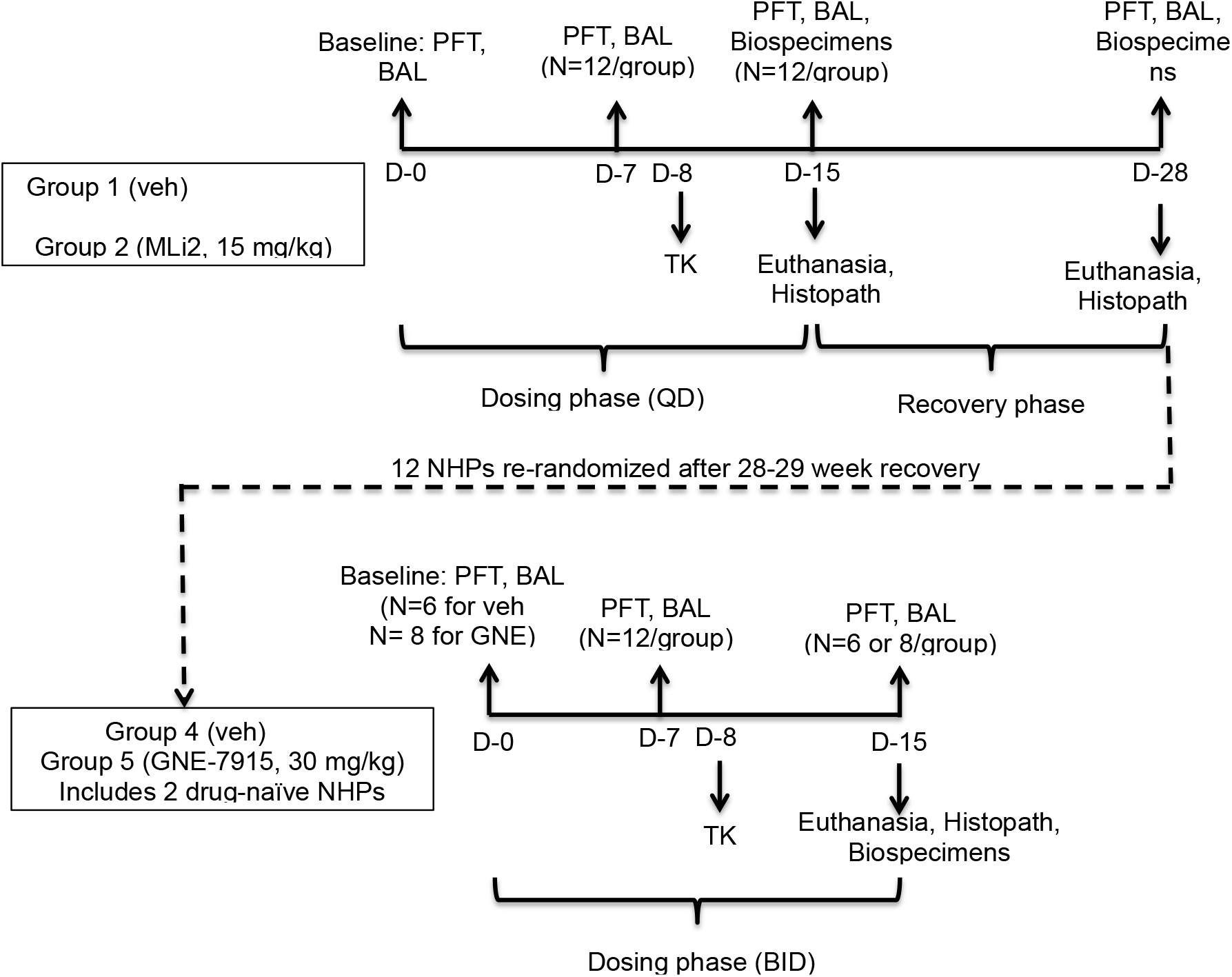
Schematic of the overall study design to assess effects on pulmonary function

#### Animals, Treatment and Testing Regimens

Female cynomolgus macaques (Chinese origin, Charles River; age range 2.5-7.2 years at the start of the study, body weight range 3.5-4.5 kg) were randomized into 3 groups of 12 individuals each to balance the body weight distribution. Group A: Vehicle (0.5% (w/v) methylcellulose); Group B: MLi-2 (15 mg/kg, PO, QD), Group C: MLi-2 (50 mg/kg, PO, QD). These groups were staggered with respect to treatments and assessments. Two animals were maintained throughout the study without any treatment and used later for GNE-7915 assessment (see below).

##### MLi-2 assessment

The test article or vehicle was administered by oral gavage once daily, in the morning, for 15 consecutive days. Following day 15 pulmonary functional tests (PFTs; see below) and biospecimen collection, a subset of non-human primates (4 from each cohort) were euthanized and submitted for necropsy and sample collection (detailed below). Another subset of 4 non-human primates went through the recovery phase (13 days) followed by euthanasia, necropsy, and biospecimen collection.

##### GNE-7915 assessment

The remaining 4 animals from each treatment group of the MLi-2 study (total N =12) and two naïve animals from the original cohort were allowed to recover for an additional 28-29 weeks. These NHPs were re-randomized into two groups, considering their previous treatment allocation in order to evenly distribute them into vehicle (n=6) and GNE-7915 (n=8) treatment groups. Vehicle and GNE-7915 (30 mg/kg) were administered twice daily, about 8 hours apart, by oral gavage for 15 consecutive days. Immediately after last PFT measurements and BAL/blood collections, all animals were euthanized and submitted for necropsy and sample collection to determine any effects of treatment.

#### Biospecimen collection

Blood was collected by venipuncture for TK and peripheral blood mononuclear cells (PBMC) isolation. Blood (< 2 mL) was collected into tubes containing EDTA. For MLi-2, blood was collected prior to dosing, and 0.5, 1, 4, 8 hours post-dose on days 1, 8, 15 and 28 prior to PFT assessments and BAL. For GNE-7915, blood samples for TK analysis were collected prior to dosing, and 0.5, 1, 4 and 8 hours post-dose on days 1 and 8 of treatment. Additional blood samples were taken on day 15 prior to PFT and processed for PBMC isolation and clinical chemistry.

#### Anesthesia and Pulmonary Function Tests (PFT)

PFTs were administered at baseline (day 0) and on days 7, 15 and 28 (for MLi-2 only). Animals were administered ketamine (5-10 mg/kg, IM) followed by isoflurane (5% for induction, 1.5-3% for maintenance). Lung diffusion capacity for carbon monoxide (DLCO), quasi-static lung compliance, forced maneuver ventilation capacities to include forced vital capacity, forced expiratory volume, forced expiratory flows, bronchoalveolar lavage of two lung segments were conducted. Animals were anesthetized and intubated with an endotracheal tube of appropriate size for the animal. The animal was placed in a heated whole-body volume displacement plethysmograph.

DLCO, quasistatic lung compliance (Cqs), and forced expiratory flow-volume curves, including forced expiratory volume (FEV) were measured during induced inhalations, exhalations during transient apnea, and then inflating and deflating the lung using either a syringe or an automated system of valves and pressure reservoirs calibrated for this purpose.

##### DLCO

The single-breath method was used to measure DLCO. The inflation volume required to reach +20 cm H2O Ptp was determined prior to each measurement. A syringe was flushed and filled with that volume of test gas (mixture of CO and Ne in air). The lung was insufflated with the test gas and a period of “breath-hold” at end inspiration (6s) occurred followed by withdrawal of 75% of the insufflated volume. A portion of the gas remaining in the airway was withdrawn into a small gas-tight syringe, and its composition was analyzed using gas chromatography. The inhaled and exhaled gas concentrations, inflation volume, and inflation time were recorded and used to calculate DLCO, which is the pressure-adjusted uptake rate of CO across the alveolar-capillary membrane.

##### Quasi-static lung compliance (Cqs10)

The Cqs maneuver was performed by insufflating the lungs slowly to total lung capacity, which was defined as the lung volume at a Ptp of +30 cm H2O, and then deflating the lung slowly until flow stopped. Quasistatic lung compliance was calculated as the slope of the pressure-volume curve between 0 and +10 cm H2O Ptp and termed Cqs10.

##### Forced Vital Capacity (FVC), Forced Expiratory Volume (FEV), and Forced Expiratory Flow Rates (FEF)

The forced expiratory maneuver was induced during apnea by inflating the lung to total lung capacity and then rapidly deflating by opening a large bore valve to a negative pressure reservoir while recording flow, volume, and Ptp. The lung volume obtained from the forced expiratory maneuver from +30cm H2O to the point at which expiratory flow ceases is the FVC. The lung volume obtained at a given time interval, e.g. 0.1s, from the start of expiration is the FEV for that time interval. FEV at the point of maximal flow is also included. The expiratory flow rates at 75% and 25% of FVC were averaged to derive Mean Mid-Expiratory Flow Rate (MMEF). Further, the flow rates at various steps of lung volume are also presented, i.e. 75%, 50%, 25%, and 10%. The flows are also normalized to lung volume by dividing by FVC from a given maneuver.

#### Bronchoalveolar Lavage

Following pulmonary function assessment, a bronchoscope (BF-XP40, Olympus America Inc., Melville NY) was maneuvered through the endotracheal tube and wedged in approximately a 4^th^-6^th^ generation airway in the left diaphragmatic lung lobe. Three aliquots of 5ml sterile USP-grade saline were instilled and aspirated in succession followed by further aspiration with an empty syringe to recover as much as possible. The process was then repeated for the contralateral lung. Each sample was stored on ice until processing.

#### Necropsy and Pathology

Animals were euthanatized and submitted for necropsy following the 15-day treatment period (MLi-2 and GNE-7915) and the 13-day recovery period (MLi-2 only). Standard necropsies were performed in addition to collection of blood, urine and cerebrospinal fluid (CSF). Approximately half of the lung, kidney, and brain were collected and stored in fixative as described in Part 1 for histopathology and the other half was flash frozen and stored at -70° to -90°C for compound levels and biomarker analyses.

#### Statistical Analysis

Data are presented as mean and SEM for continuous variables. PFT data are analyzed using the 2-way ANOVA. Bonferroni post tests were conducted by comparing treated groups to vehicle treated group per study day and comparing each post-treatment value to corresponding pre-treatment value for a given group. A p-value of ≤0.05 indicates statistical significance. Analyses were completed using GraphPad Prism (GraphPad Software Inc., San Diego, CA, USA).

## Acknowledgments

I would like to thank Emily Murphy (MJFF) and Scott Meola (Simplissimus) for technical assistance on the manuscript preparation.

MASB, KM, BKF, TBS wrote the manuscript, designed the studies, and analyzed data. TB, DKB, ME, AAS, MJF, RNF, PG, AGH, SH, Wh, CH, MEK, XL, MLM, CM, HM, WAM, SP, CR, KR, AKS, AS, SS, CT, ZY, HY, XW designed studies, conducted experiments, and analyzed data.

---

**Supplemental Table 1:**
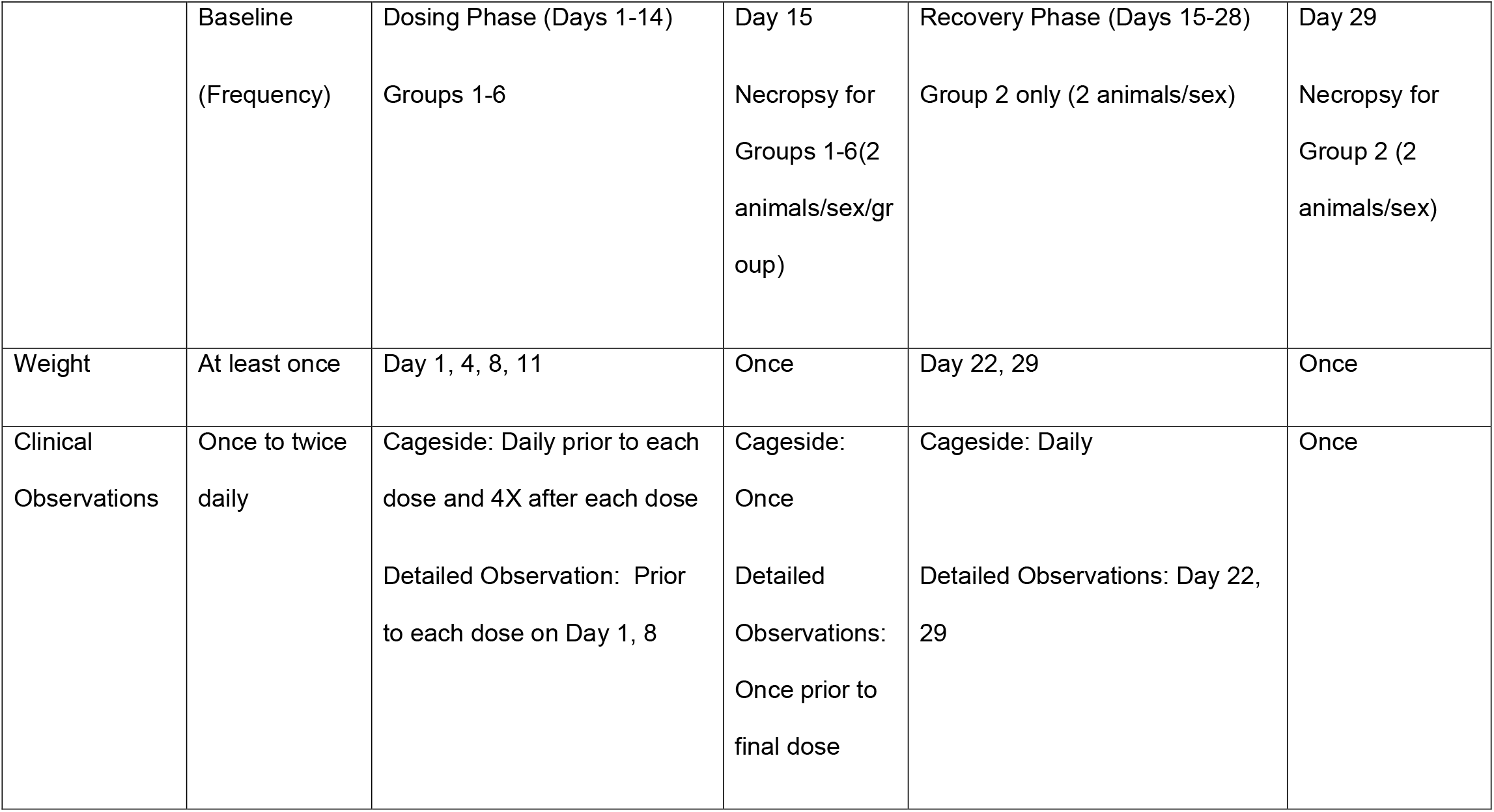
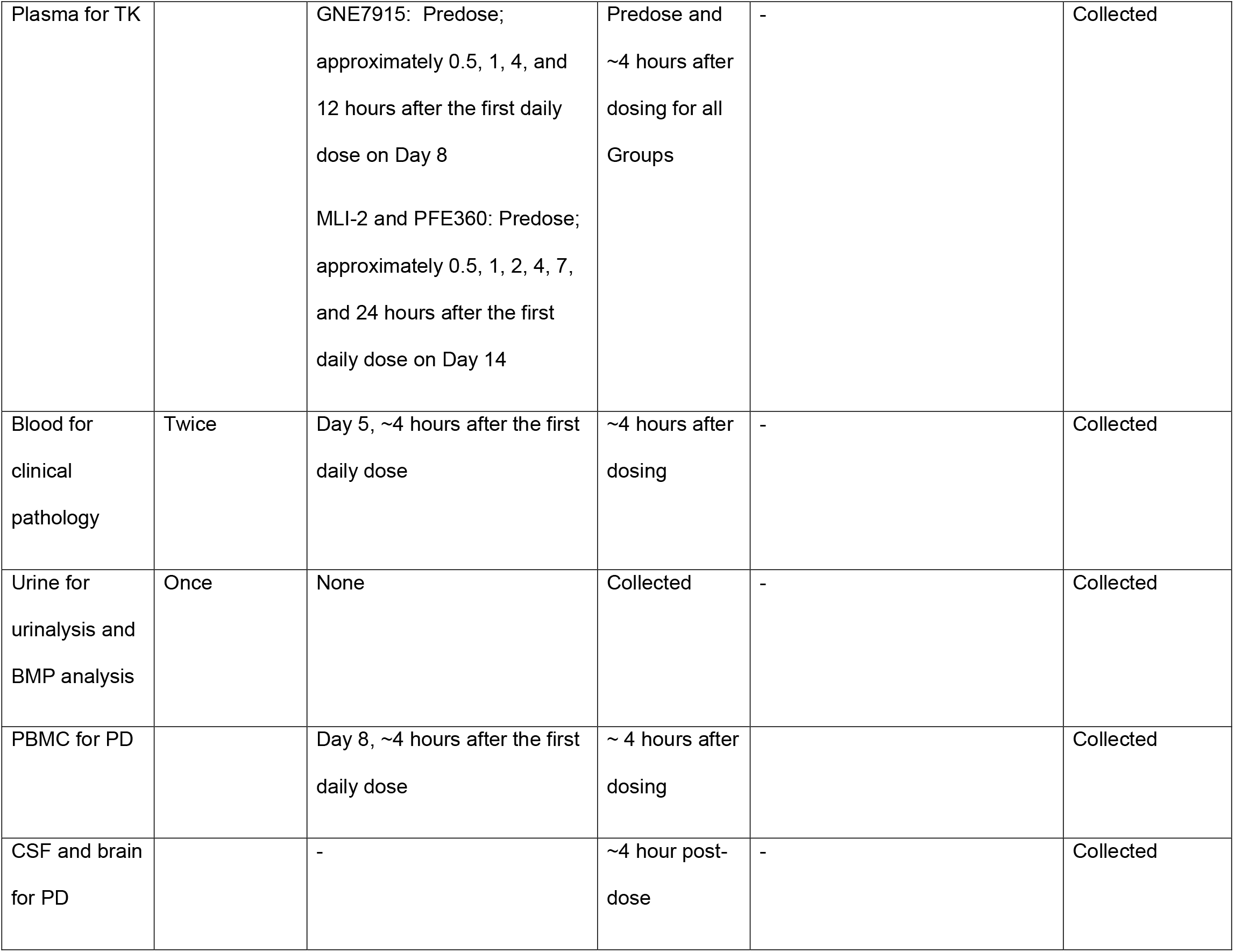
The schedule of dosing, monitoring, necropsies and tissue collections of non-human primates in Part 1

|  | Baseline<br>(Frequency) | Dosing Phase (Days 1-14)<br>Groups 1-6 | Day 15<br>Necropsy for<br>Groups 1-6(2<br>animals/sex/gr<br>oup) | Recovery Phase (Days 15-28)<br>Group 2 only (2 animals/sex) | Day 29<br>Necropsy for<br>Group 2 (2<br>animals/sex) |
| --- | --- | --- | --- | --- | --- |
| Weight | At least once | Day 1, 4, 8, 11 | Once | Day 22, 29 | Once |
| Clinical<br>Observations | Once to twice<br>daily | Cageside: Daily prior to each<br>dose and 4X after each dose<br>Detailed Observation: Prior<br>to each dose on Day 1, 8 | Cageside:<br>Once<br>Detailed<br>Observations:<br>Once prior to<br>final dose | Cageside: Daily<br>Detailed Observations: Day 22,<br>29 | Once |
| Plasma for TK |  | GNE7915: Predose;<br>approximately 0.5, 1, 4, and<br>12 hours after the first daily<br>dose on Day 8<br>MLI-2 and PFE360: Predose;<br>approximately 0.5, 1, 2, 4, 7,<br>and 24 hours after the first<br>daily dose on Day 14 | Predose and<br>~4 hours after<br>dosing for all<br>Groups | - | Collected |
| Blood for<br>clinical<br>pathology | Twice | Day 5, ~4 hours after the first<br>daily dose | ~4 hours after<br>dosing | - | Collected |
| Urine for<br>urinalysis and<br>BMP analysis | Once | None | Collected | - | Collected |
| PBMC for PD |  | Day 8, ~4 hours after the first daily dose | ~ 4 hours after dosing |  | Collected |
| CSF and brain for PD |  | - | ~4 hour post-dose | - | Collected |

